# Reorganization of the Mitochondria-Organelle Interactome during Postnatal Development in Skeletal Muscle

**DOI:** 10.1101/2021.06.16.448433

**Authors:** Yuho Kim, Eric Lindberg, Christopher K. E. Bleck, Brian Glancy

## Abstract

Cellular development requires the integrated assembly of intracellular structures into functionally specialized regions supporting overall cellular performance. However, it remains unclear how coordination of organelle interactions contributes to development of functional specificity across cell types. Here, we utilize a subcellular connectomics approach to define the cell-scale reorganization of the mitochondria-organelle interactome across postnatal development in skeletal muscle. We show that while mitochondrial networks are disorganized and loosely associated with the contractile apparatus at birth, contact sites among mitochondria, lipid droplets, and the sarcoplasmic reticulum are highly abundant in neonatal muscles. The maturation process is characterized by a transition to highly organized mitochondrial networks wrapped tightly around the muscle sarcomere but also to less frequent interactions with both lipid droplets and the sarcoplasmic reticulum. These data demonstrate a developmental redesign reflecting a functional shift from muscle cell assembly supported by inter-organelle communication toward a muscle fiber highly specialized for contractile function.

## Introduction

Cellular assembly necessitates the physical coordination of many different organelles in order to optimize intracellular structure to meet the functional requirements of the cell^1,2^. Thus, with the extensive array of functional demands observed across cell types within the body, the internal structure within different cells can also vary widely^3,4^. While the functional consequences of altering the content of different organelle types within a cell is well appreciated^4^, and the significance of varying organelle protein and lipid composition is increasingly recognized^5,6^, the impact of organelle organization on overall cellular function is less well understood^7^. Organelles within a cell do not operate in isolation, but rather they rely on inputs from and/or interactions with other cellular components in order to perform tasks in support of the cell^1,7^. As such, the spatial proximity among different organelles determines how quickly or how often interactions can occur, both of which are critical regulators of the functional capacity for a given process within a cell. Thus, a better understanding of how organelle interactions are altered in response to changing functional demands would provide key insight into how intracellular organization contributes to cellular function. Additionally, while the impact of organelle interactions in cell culture^8,9^, adult tissues^5,10^, and pathological conditions^11,12^ has been of great interest in recent years, there is little information available on the role of interorganelle connectivity during cellular assembly or development when communication and coordination among organelles is likely critical.

Mitochondria are extensively associated with other organelles as part of the cellular energy distribution system^1,13^ and these interactions are crucial for cellular metabolism and function^1,14^. For instance, mitochondria can form direct contact sites with their lipid droplet (LD) fuel source, and these mitochondria are reported to be larger, longer, and have greater energy conversion capacity compared to non-LD connected mitochondria within the same cell^10,15,16^. Additionally, the dynamic nature of the physical contacts between mitochondria and the endo/sarcoplasmic reticulum^17,18^ allows for firm regulation over both mitochondrial and cytosolic calcium levels which play a critical role in many cellular processes including energy homeostasis and cell viability^19-21^. In striated muscle cells, mitochondria can also be closely associated with the high energy demanding contractile machinery which takes up the majority of cellular volume. However, it remains unclear how mitochondria in different cell types and/or physiologic environments balance the need for physical interactions with multiple cellular structures as well as the cytosol in support of overall cellular function.

Here, we define the physical reorganization of the cellular energy distribution system supporting sustained skeletal muscle contraction across postnatal development. By combining the large field of view and nanometer resolution afforded by focused ion beam scanning electron microscopy (FIB-SEM)^22,23^ with machine learning image segmentation^24^, we provide a high-throughput subcellular connectomics analysis^10^ of the 3D mitochondria-organelle interactions within developing skeletal muscle. We find that while tortuous mitochondria are loosely interspersed within the contractile machinery at birth^25,26^, frequent interactions among mitochondria, lipid droplets, and the sarcoplasmic reticulum take place in neonatal muscles. During maturation into either oxidative or glycolytic muscle types, a structural transition occurs where mitochondria become more linear in nature as well as more tightly associated with the contractile apparatus^25,26^. However, muscle maturation is also characterized by less frequent interactions among mitochondria, lipid droplets, and sarcoplasmic reticulum including a near complete loss of lipid droplets in glycolytic muscle. These results reflect a functional redesign of the skeletal muscle cell during postnatal development where frequent organelle interactions support the need for coordinated cellular assembly at birth while the cellular energy distribution system becomes specifically tuned to support contractile function in the mature muscle cell.

## Results

### Dynamic reorganization of mitochondrial networks during postnatal development

To evaluate the mitochondria-organelle interactome during postnatal muscle development, we used FIB-SEM to collect mouse muscle cell volumes with 10 nm resolution in 3D at birth (postnatal day 1 (P1), **Supplementary Movie 1**), during the dynamic phase of the transition between neonatal and mature mitochondrial networks^27^ (P14, **Supplementary Movie 2**), and after mitochondrial network structures had reached maturity^27^ (P42, **Supplementary Movies 3-4**). To account for the muscle fiber type differences in mitochondrial intra- and inter-organelle interactions previously observed in mature muscles^10^, we imaged cells from both the soleus and gastrocnemius muscles representing more oxidative and glycolytic muscles, respectively^28^, and further confirmed cell type based on mitochondrial content^10^. Machine learning segmentation of the FIB-SEM muscle volumes^10,24,29^ allowed for high-throughput analyses of mitochondrial, lipid droplet (LD), and sarcotubular (SR/T) structures as well as interactions among them.

Mitochondrial structure within a cell is coordinated across different spatial scales ranging from cell-wide networks to the size and shape of individual organelles to interactions with adjacent organelles^3,10^ (**Figure 1a-f**). Beginning at the cellular scale, networks of tortuous mitochondria were primarily aligned parallel to but loosely associated with the contractile apparatus in both newborn muscle types (**Figure 1a,d,I**, **Supplementary Movie 1**). Additionally, overall mitochondrial volume and number were similar between muscle types at birth (**Figure 1g-h**, 6.3±1.2% and 7.0±0.4% of total muscle volume, 345±20 and 313±17 mitochondria/1000 µm^3^ muscle, mean±SE, n=3 muscle volumes, 618 and 276 mitochondria for P1 soleus and gastrocnemius, respectively). These data suggest that mitochondrial network configuration in neonatal muscles is driven by developmental status rather than muscle type at this stage. During the postnatal transition phase (P14), overall mitochondrial content and number were little changed from birth and were again no different between muscle types (**Figure 1g-h**, 10.8±0.2% and 7.7±1.9% of total muscle volume, 400±26 and 443±6 mitochondria/1000 µm^3^ muscle, mean±SE, n=3 muscle volumes, 1124 and 1848 mitochondria for P14 soleus and gastrocnemius, respectively). However, mitochondrial networks in both muscles began to more closely associate with the contractile apparatus at P14 (**Figure 1b,e**, **Supplementary Movie 2**). In the soleus, mitochondrial networks became more linear and elongated along the parallel axis and began to form short branches along the perpendicular axis at the ends of the sarcomeres near the z-disk (**Figure 1b,i**). In contrast, while the mitochondrial networks in the gastrocnemius muscle also became more linear and elongated compared to at birth (**Figure 1e**), there was a greater contribution of perpendicularly oriented mitochondrial branches compared to the soleus muscle (**Figure 1i**). By P42, the divergence of mitochondrial network configurations between muscle types was completed (**Figure 1c,f**) as mitochondrial volume and number were significantly higher in the oxidative compared to glycolytic muscles (**Figure 1g-h**, 10.8±2.2% and 3.6±0.3% of total muscle volume, 420±37 and 247±25 mitochondria/1000 µm^3^ muscle, mean±SE, n=3 muscle volumes, 1147 and 475 mitochondria for P42 oxidative and glycolytic muscles, respectively), and the mitochondrial networks reached their mature grid-like (oxidative, **Supplementary Movie 3**) and primarily perpendicular (glycolytic, **Supplementary Movie 4**) orientations (**Figure 1c,f,i**). These data suggest that the timing of muscle fiber-type specificity of mitochondrial network structure occurs in concert with the fiber-type specificity of myosin isoform composition that occurs during postnatal development^30,31^.

**Figure 1:**
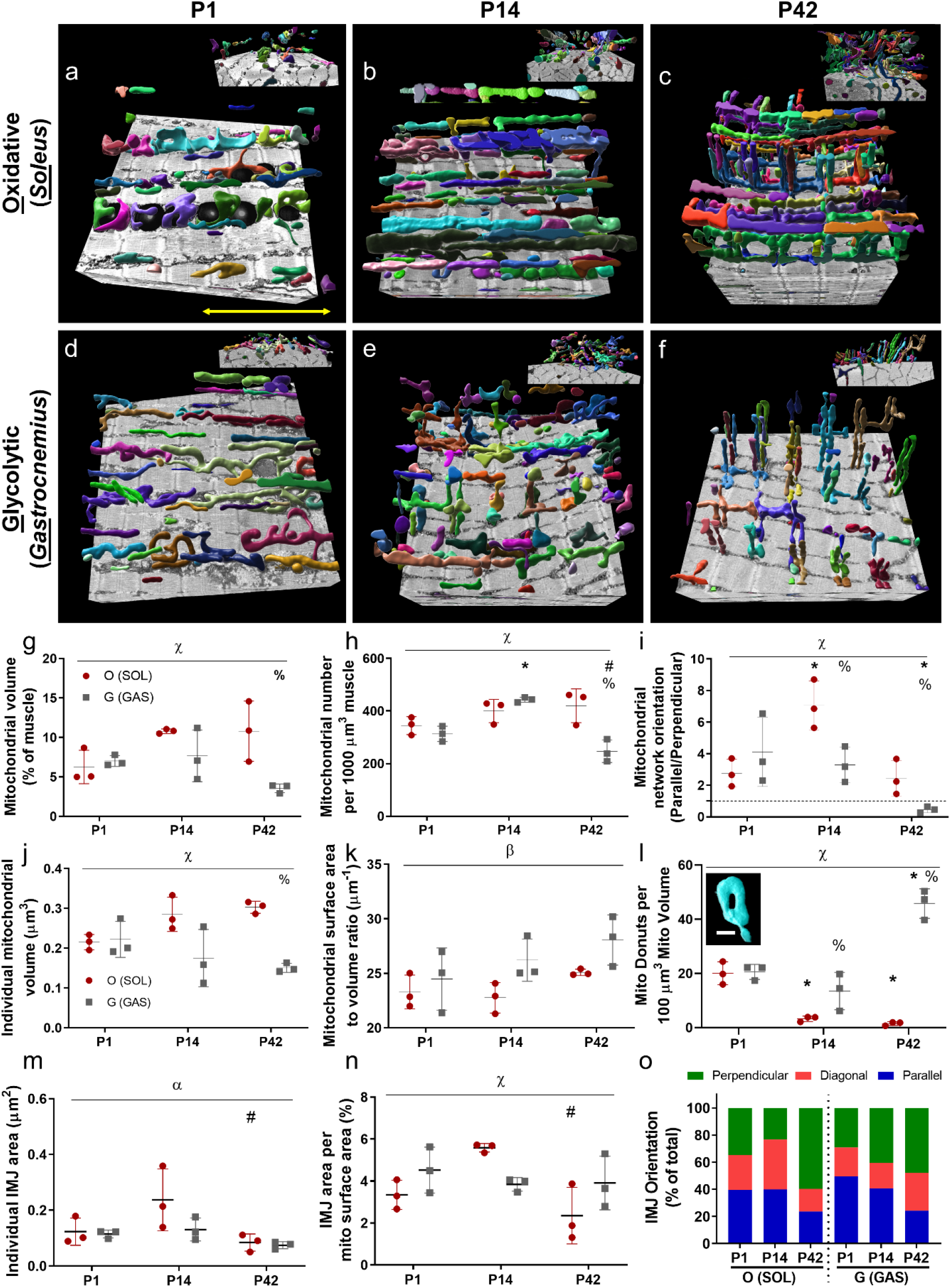
Dynamic reorganization of mitochondrial networks during postnatal development. **a-f)** Representative 3D rendering of individual mitochondria within networks in oxidative (O; SOL) and glycolytic (G; GAS) fibers of mice at postnatal (P) day 1, 14, and 42, respectively. Mitochondrial networks are arranged along muscle contraction axis (yellow arrow) and 90-degree rotated images are depicted in the upper-right corner. Each color indicates individual mitochondria. **g)** Total mitochondrial volume (% of muscle area). **h)** Mitochondrial number per 1000 μm^3^ of muscle. **i)** Mitochondrial network orientations are calculated in ratio of parallel to perpendicular arrangement. **j)** Individual mitochondrial volume (μm^3^). **k)** Mitochondrial surface area to volume ratio (μm^-1^). **l)** Donut-shaped mitochondria are counted per 100 μm^3^ of mitochondrial volume. Representative image is displayed in the upper-right corner. **m)** Individual IMJ area (μm^3^). **n)** IMJ area per mitochondrial surface area (%). **o)** Quantification of intermitochondrial junction (IMJ) orientation. N values: P1 oxidative – 609 mitochondria, 375 IMJ, 3 datasets; P14 oxidative – 1115 mitochondria, 523 IMJ, 3 datasets; P42 oxidative – 1414 mitochondria, 509 IMJ, 3 datasets; P1 glycolytic – 274 mitochondria, 173 IMJ, 3 datasets; P14 glycolytic – 1828 mitochondria, 786 IMJ, 3 datasets; P42 glycolytic – 462 mitochondria, 263 IMJ, 3 datasets. Points are means for each dataset. Bars are means ± SE. \**P < 0*.*05*, vs P1; ^#^*P < 0*.*05*, vs P14; ^%^*P < 0*.*05*, vs O (SOL);; ^α^*P<0*.*05*, main effect of development; ^β^*P<0*.*05*, main effect of fiber type; ^χ^*P<0*.*05*, interaction effect of development and fiber type. scale bar = 1 μm.

At the single organelle level, the volume of individual mitochondria followed a similar time course to the overall mitochondrial content in both muscle types where content gradually increased across development in the oxidative muscles and fell in the glycolytic muscles (**Figure 1j**). These data indicate that mitochondrial functional capacity across both the network and individual organelle level may be coordinated together^3^. While there were no significant differences in mitochondrial surface area to volume (SA/V) ratio among individual groups (**Figure 1k**), gastrocnemius muscle mitochondria as a whole had greater SA/V ratios compared to soleus muscle mitochondria consistent with previous reports on glycolytic versus oxidative muscle mitochondria^10,32,33^. To further investigate how individual muscle mitochondrial morphology is altered across postnatal development, we quantified the relative number of small (∼80-120 nm) donut-like holes in mitochondria (**Figure 1l inset**) which have been suggested as a marker of oxidative stress^34^. In soleus muscles, the number of mitochondrial donuts was highest at birth then dropped significantly at P14 and remained low in the mature muscle (**Figure 1l**, 20.0±2.4, 3.3±0.6, and 1.5±0.4 mito donuts per 100 µm^3^ mito volume, n=3 muscle volumes, 26, 10, and 5 donuts for P1, P14, and P42, respectively). In the gastrocnemius, there were no differences in the number of donuts compared to the soleus at birth. However, the relative number of mitochondrial donuts remained elevated at P14 and rose significantly to more than thirty-fold higher than in the oxidative muscles at P42 (**Figure 1l**, 20.6±1.6, 13.5±4.0, and 45.8±3.1 mito donuts per 100 µm^3^ mito volume, n=3 muscle volumes, 17, 24, and 32 donuts for P1, P14, and P42, respectively), suggesting that the increased oxidative stress reported previously in glycolytic muscles^10^ may be reflected at the mitochondrial level beginning during the late postnatal phase of development.

To determine whether interactions among mitochondria were altered during postnatal muscle development, we assessed the size, abundance, and orientation of the intermitochondrial junctions (IMJs) between adjacent mitochondria which have been suggested to allow for rapid communication and distribution of molecules among physically coupled mitochondria^35^. The size of individual IMJs and relative abundance of IMJs per mitochondrion were both largely similar across muscle types and developmental timepoints with the exception of an increase in both size and abundance observed at P14 in the soleus compared to the mature oxidative muscle (**Figure 1m,n**). Conversely, while there were no differences in IMJ orientation detected between muscle types, each muscle types demonstrated a loss of parallel and a gain of perpendicularly oriented IMJs upon maturation (**Figure 1o**). These data suggest that the putative physical coupling sites permitting transfer of signaling molecules, metabolites, and/or ions directly between adjacent mitochondria are primarily altered by changing their orientation within the cell rather than size or abundance during postnatal development.

### Mitochondria-lipid droplet (LD) contact sites decrease in frequency across postnatal development

To evaluate how mitochondrial interactions with other organelles were altered during postnatal muscle development, we began by assessing the size and content of the lipid droplets (**Figure 2a-c**) which provide a direct fuel source for mitochondrial oxidative phosphorylation. Overall muscle content and the size of individual lipid droplets were highest at birth in both muscle types followed by a significant decrease during the late postnatal stage that continued into maturation (**Figure 2d,e**). Contact sites between mitochondria and lipid droplets (i.e., membranes within 30 nm^36^) followed a similar pattern with a nearly ten-fold loss in lipid droplet contact site abundance per mitochondrion across postnatal development in the oxidative muscles and a complete loss of contact sites in mature glycolytic muscle where no lipid droplets were found (**Figure 2f**). These data suggest that physical interactions between mitochondria and lipid droplets, which facilitate direct transfer of molecules between them, may be directly related to the metabolic fuel preferences of the skeletal muscle cell, as neonatal muscles are known to rely more heavily on fatty acids compared to adult muscles^37^, while glycolytic muscles rely more on carbohydrate fuel sources relative to oxidative muscles^38^.

**Figure 2:**
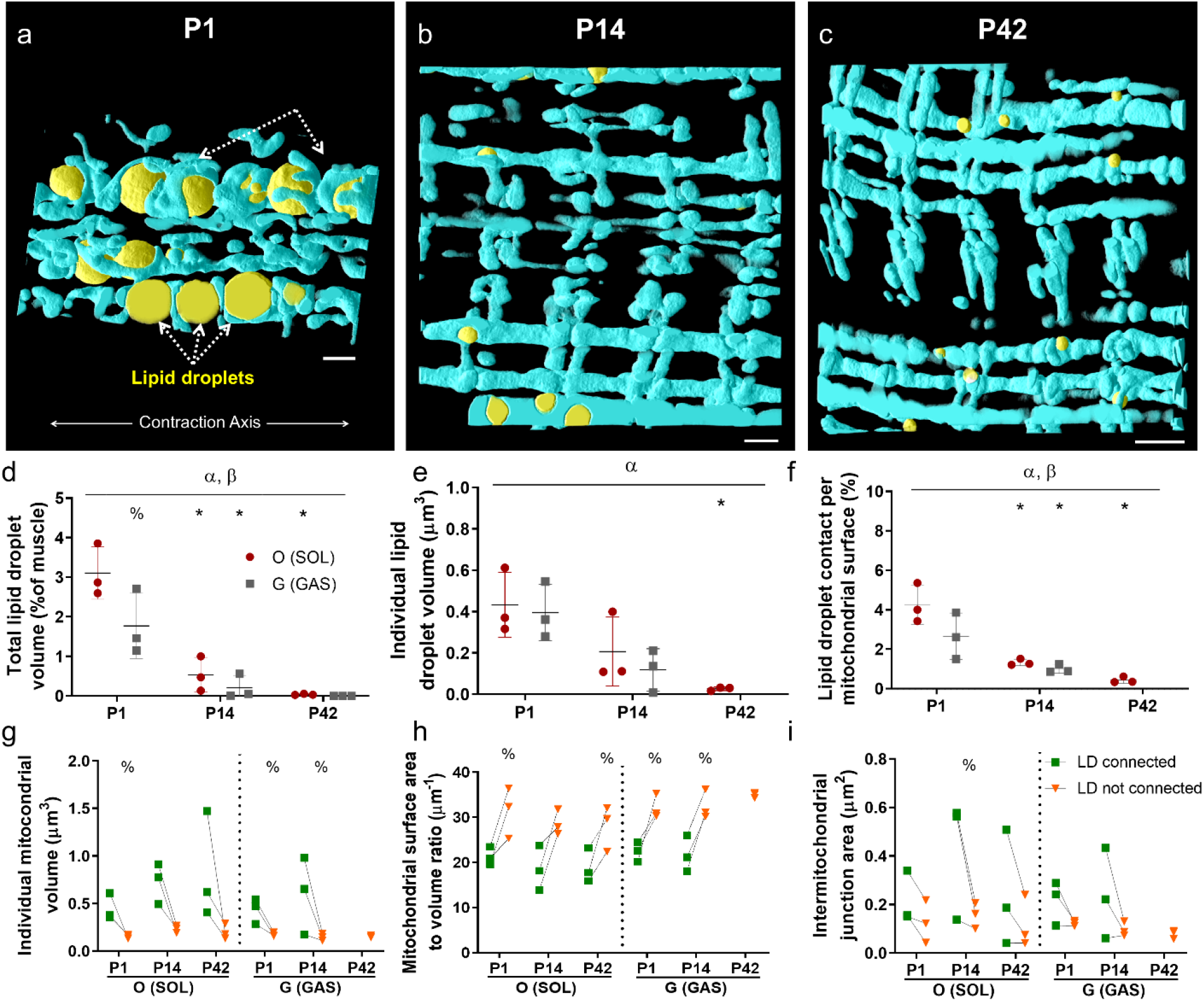
Mitochondria-lipid droplet (LD) contact sites decrease in frequency across postnatal development. **a-c)** Representative 3D rendering of mitochondrial networks (colored in sky blue) and lipid droplets (colored in yellow) of oxidative fibers of mice at postnatal (P) day 1, 14, and 42, respectively. All images are aligned to contraction axis. **d)** Total lipid droplet volume (% of muscle). **e)** Individual lipid droplet volume (μm^3^). N values: P1 oxidative – 183 LD, 3 datasets; P14 oxidative – 117 LD, 3 datasets; P42 – 81 LD, 3 datasets; P1 glycolytic – 53 LD, 3 datasets; P14 glycolytic – 89 LD, 3 datasets; P42 glycolytic – 0 LD, 3 datasets. **f)** LD contact per mitochondrial surface area (%). **g-i)** Individual mitochondrial volume (μm^3^; g), Mitochondrial surface area to volume ratio (μm^-1^; h), and Intermitochondrial junction area (μm^2^; i) connected with LD (*LD connected*) and non-connected with LD (*LD not connected*). N values (*LD connected*): P1 oxidative – 183 mitochondria, 3 datasets; P14 oxidative – 117 mitochondria, 3 datasets; P42 – 81 mitochondria, 3 datasets; P1 glycolytic – 53 mitochondria, 3 datasets; P14 glycolytic – 89 mitochondria, 3 datasets; P42 glycolytic – 0 mitochondria, 3 datasets; N values (*LD not connected*): P1 oxidative – 414 mitochondria, 3 datasets; P14 oxidative – 406 mitochondria, 3 datasets; P42 – 428 mitochondria, 3 datasets; P1 glycolytic – 219 mitochondria, 3 datasets; P14 glycolytic – 697 mitochondria, 3 datasets; P42 glycolytic – 263 mitochondria, 3 datasets. Points are means for each dataset. Bars represent means±SE. \**P < 0*.*05*, vs P1; ^%^*P < 0*.*05*, vs O (SOL); ^α^*P<0*.*05*, main effect of development. ^β^*P<0*.*05*, main effect of fiber type. Scale bar = 1 μm.

We previously found that mitochondria in contact with lipid droplets provided a structural capacity for greater energy distribution compared to non-lipid connected mitochondria within the same adult muscle cells^10^. To determine whether this apparent structural specialization of mitochondria within a network was present beginning at birth, we assessed individual mitochondrial structural characteristics for lipid droplet connected and non-connected mitochondria separately. Individual mitochondrial volume, a proxy for the internal capacity of a mitochondrion, was consistently greater in lipid connected mitochondria for all muscles with lipid droplets present (**Figure 2g**). Conversely, mitochondrial SA/V ratio, an indicator of the relative capacity to interact with the surrounding environment, was lower in lipid droplet connected mitochondria across all timepoints and in both muscle types (**Figure 2h**). Additionally, the total IMJ area per mitochondrion, an indicator of molecular transfer capacity to mitochondria, trended higher in lipid droplet connected mitochondria (**Figure 2i**). Together, these data indicate that the structural, and likely functional, specialization of lipid droplet connected mitochondria for energy distribution capacity rather than interaction capacity has already begun at birth and is maintained throughout the maturation process.

### Mitochondria-sarcoplasmic reticulum interactions are highly abundant during early postnatal development

To begin our evaluation of mitochondrial interactions with the SR/T throughout postnatal development, we first assessed the total muscle cell volume occupied by the SR/T. At birth, the SR/T formed an unorganized mesh wrapping around the myofibrils with no difference in total volume between muscle types (**Figure 3a,c,e**), similar to previous work in mice^39,40^. During maturation, the SR/T became more organized into the well-known longitudinal SR mesh and SR/T triads wrapping around the myofibrils (**Figure 3b,d**) with SR/T in the mature glycolytic muscle occupying a greater volume than in newborn muscles or mature oxidative muscle (**Figure 3e**), also in line with previous reports^10,39,41^. These data demonstrate the cell-type specification of the SR/T as well as the consistency between our 3D analysis and previous 2D assessments of SR/T volume.

Next, we assessed the physical interactions between the SR/T and 5332 individual mitochondria during postnatal development. Nearly, every mitochondrion across all conditions had at least one contact (membranes within 30 nm) with the SR/T (99.6±0.5, 99.6±0.2, 99.4±0.2, 99.3±0.3, 99.6±0.2, 98.3±1.2% of mitochondria in contact with SR/T, n=300, 570, 1847, 1124, 468, 1023 mitochondria in 3 datasets for P1 gastrocnemius, P1 soleus, P14 gastrocnemius, P14 soleus, P42 glycolytic, and P42 oxidative, respectively). The triadic nature of the interactions between the t-tubules and the sarcoplasmic reticulum (SR) means that most of the t-tubule surface is covered by the SR^40^, and, as a result, the mitochondrial interactions with the SR/T volume detected here are almost exclusively interactions between mitochondria and the SR. At birth, roughly one-sixth of the mitochondrial surface area was in direct contact with the SR/T on average with more than 80% of mitochondria having at least 10% of its surface area in contact with the SR/T (**Figure 4f,g**, 16.6±2.2 and 17.3±2.3% mitochondrial surface area contact with SR/T, 83.5±7.9 and 84.3±10.7% of mitochondria with >10% surface area contact with SR/T, n=300 and 570 mitochondria in 3 datasets for P1 gastrocnemius and P1 soleus, respectively). By two weeks of age, interactions between mitochondria and the SR/T fell by more than two-fold for mean surface area contact and more than four-fold for percentage of mitochondria with at least 10% surface area in contact and were maintained at this level into maturity for both muscle types (**Figure 4f,g**, 7.0±1.1, 6.1±0.6, 8.1±0.6, and 6.8±0.4% mitochondrial surface area contact with SR/T, 18.0±6.7, 15.2±1.9, 18.5±1.1, and 17.1±1.8% of mitochondria with >10% surface area contact with SR/T, n=1847, 1124, 468, and 1023 mitochondria in 3 datasets for P14 gastrocnemius, P14 soleus, P42 glycolytic and P42 oxidative, respectively). These data demonstrate the ubiquitous yet dynamic nature of mitochondrial interactions with the SR in muscle cells and suggests that mitochondria with high SR/T contact areas may be tailored for a different functional specialization compared with lower SR/T contacting mitochondria.

**Figure 3:**
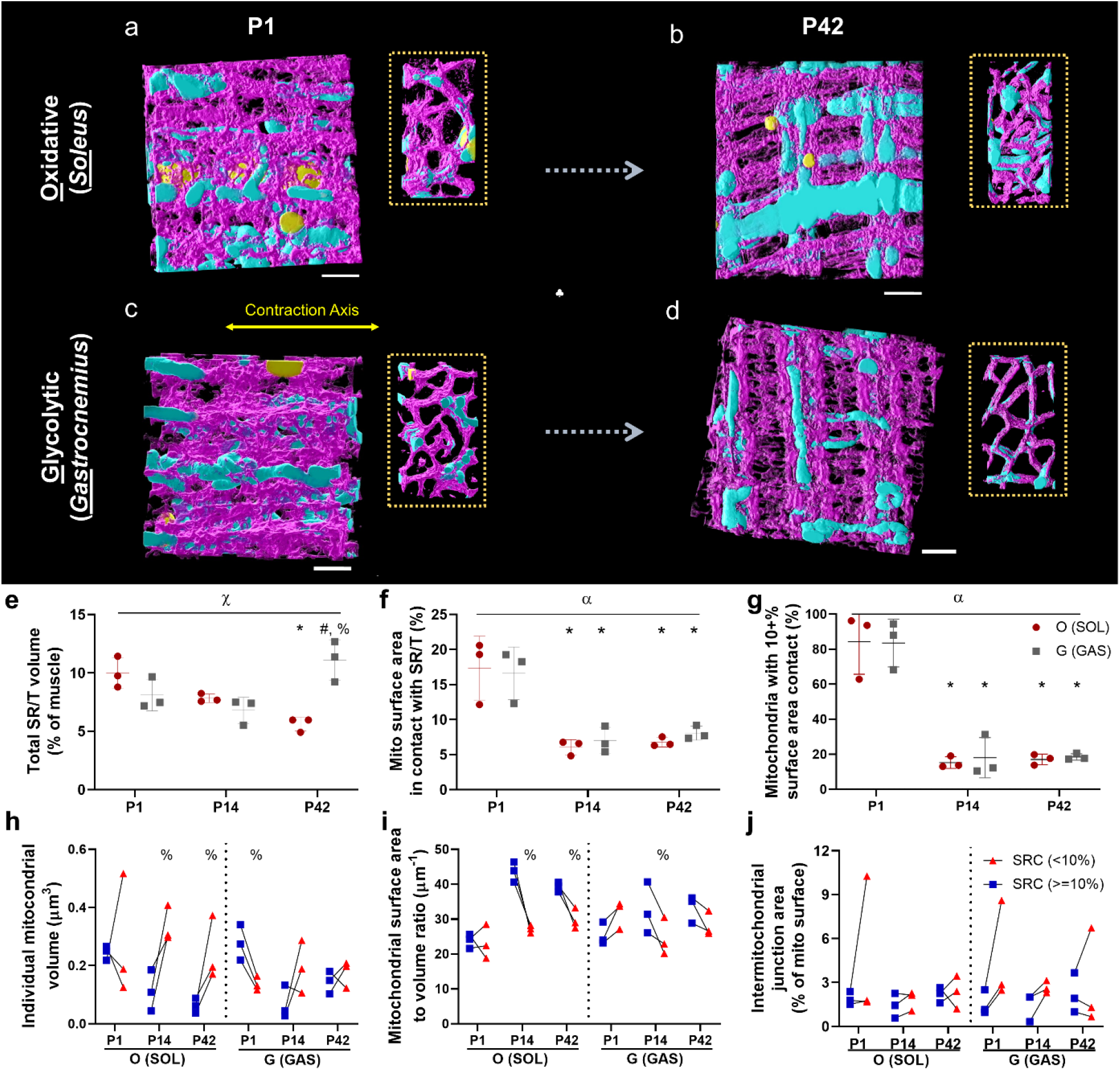
Mitochondria-sarcoplasmic reticulum interactions are highly abundant during early postnatal development. **a-d)** Representative 3D rendering of mitochondrial network (colored in sky blue color), sarcoplasmic reticulum/t-tubules (SR/T; colored in magenta), and lipid droplets (colored in yellow) in oxidative (O: SOL) and glycolytic (G: GAS) fibers of mice at postnatal (P) day 1 and 42, respectively. These 3D images are arranged along muscle contraction axis and the 90-degree rotated images are depicted in the box of dotted lines. **e)** Total SR/T volume (% of muscle). **f)** Mitochondrial surface area in contact with SR/T (%). N values: P1 oxidative – 566 Mitochondria in contact with SR/T (M-SR/T), 3 datasets; P14 oxidative – 1111 M-SR/T, 3 datasets; P42 – 1023 M-SR/T, 3 datasets; P1 glycolytic – 296 M-SR/T, 3 datasets; P14 glycolytic – 1811 M-SR/T, 3 datasets; P42 glycolytic – 458 M-SR/T, 3 datasets. **g)** Mitochondria with at least 10% surface area contact with SR/T (%). N values: P1 oxidative – 566 Mitochondria in contact with SR/T (M-SR/T), 3 datasets; P14 oxidative – 1111 M-SR/T, 3 datasets; P42 – 1023 M-SR/T, 3 datasets; P1 glycolytic – 296 M-SR/T, 3 datasets; P14 glycolytic – 1811 M-SR/T, 3 datasets; P42 glycolytic – 458 M-SR/T, 3 datasets **h-j)** Individual mitochondrial volume (μm^3^; h), Mitochondrial surface area to volume ratio (μm^-1^; i), and Intermitochondrial junction area (% of mito surface area; j) in mitochondria highly connected with SR/T (*>=10% SRC* [large area connected] vs. *<10% SRC* [less area connected]). N values (*>=10% SRC*): P1 oxidative – 494 mitochondria, 3 datasets; P14 oxidative – 131 mitochondria, 3 datasets; P42 – 174 mitochondria, 3 datasets; P1 glycolytic – 252 mitochondria, 3 datasets; P14 glycolytic – 354 mitochondria, 3 datasets; P42 glycolytic – 110 mitochondria, 3 datasets; N values (*<10% SRC*): P1 oxidative – 72 mitochondria, 3 datasets; P14 oxidative – 980 mitochondria, 3 datasets; P42 – 849 mitochondria, 3 datasets; P1 glycolytic – 44 mitochondria, 3 datasets; P14 glycolytic – 1457 mitochondria, 3 datasets; P42 glycolytic – 348 mitochondria, 3 datasets. Points are means for each dataset. Bars represent means±SE. \**P < 0*.*05*, vs P1; ^%^*P < 0*.*05*, vs O (SOL); ^#^*P < 0*.*05*, vs P14; ^χ^*P < 0*.*05*, interaction effect of development and fiber type; ^α^*P<0*.*05*, main effect of development. Scale bar = 1 μm.

**Figure 4:**
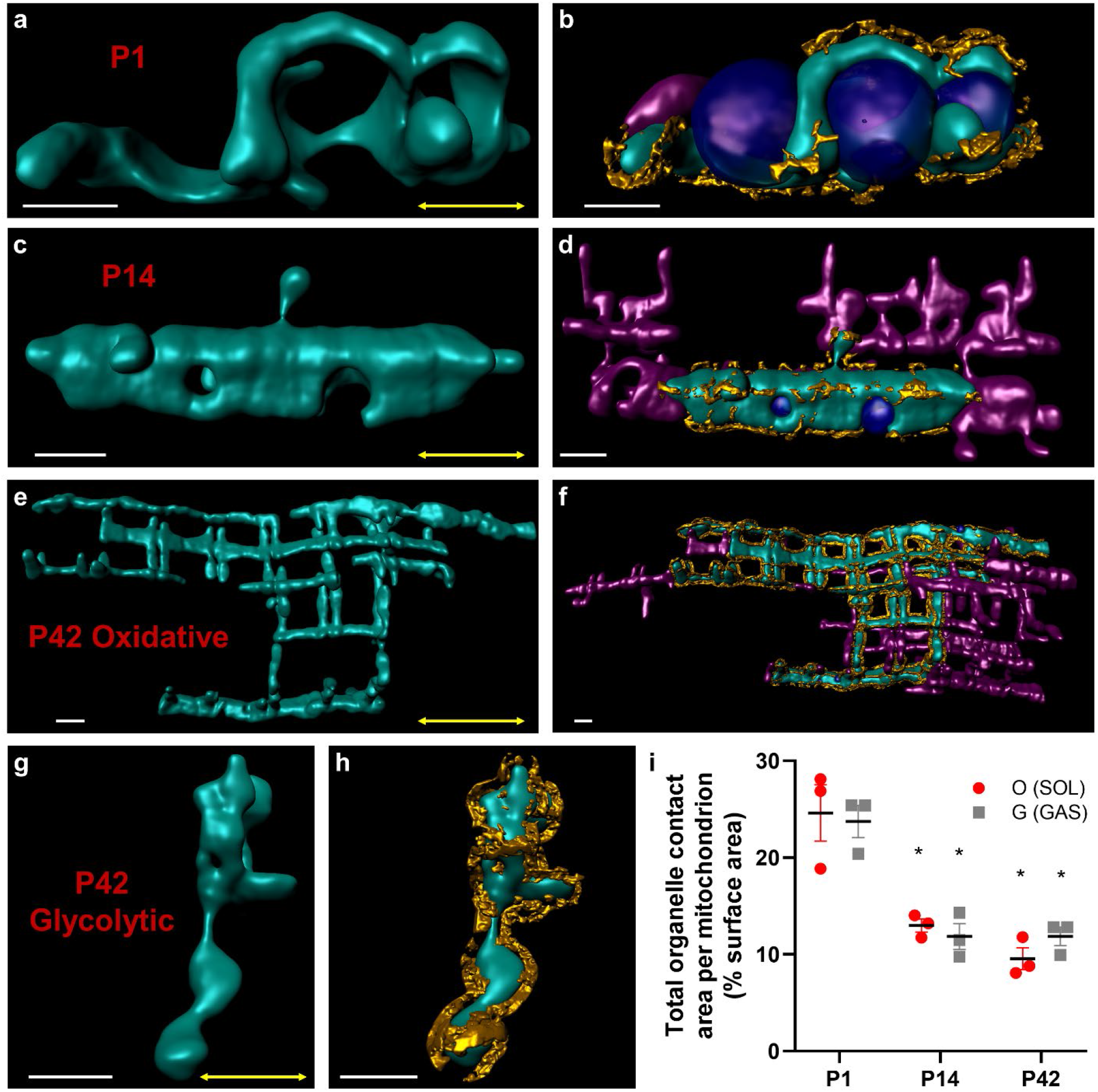
Specialized mitochondria-organelle interactions transition from supporting cellular assembly to contractile function in skeletal muscle cells. **a,b)** Representative single P1 mitochondrion (green) alone (a) and in contact (b) with other mitochondria (magenta), lipid droplets (blue), and SR/T (gold). **c,d)** Representative single P14 mitochondrion alone (c) and in contact (d) with other mitochondria, lipid droplets, and SR/T. **e**,**f)** Representative single P42 oxidative mitochondrion alone (e) and in contact (f) with other mitochondria, lipid droplets, and SR/T. **g**,**h)** Representative single P42 glycolytic mitochondrion alone (g) and in contact (h) with the SR/T. **i)** Total mitochondria-organelle contact area (%) per mitochondrion. Data equivalent to sum of figures 1n+2f+3f.

To investigate a potential difference in functional specialization among mitochondria with high and low SR/T contact, we compared the structural capacities of mitochondria with more or less than 10% surface area contact with the SR/T (**Figure 4g-j**). At birth, high SR/T contact mitochondria were larger and had lower SA/V ratios than low SR/T contact mitochondria in gastrocnemius muscle (**Figure 4h,i**), suggesting a relatively greater internal capacity but lower interaction capacity for high SR/T contact mitochondria. However, this trend was reversed by P14 and into maturity where high SR/T contact mitochondria were smaller and had higher SA/V ratios compared to low SR/T contact mitochondria in soleus muscles (**Figure 4h,i**). There were no significant differences in mitochondrion-to-mitochondrion interactions through IMJs between high and low SR/T contact groups (**Figure 4j**). These data imply that the functional specificity of mitochondria-SR membrane contact sites may undergo a developmental switch and are not directly related to the capacity for molecular transfer through mitochondrial networks.

### Specialized mitochondria-organelle interactions are a key feature of cellular assembly during postnatal development

To evaluate the integrated role of mitochondria-organelle contact sites during postnatal muscle development, we assessed how the total mitochondrial outer membrane surface area dedicated to membrane contact sites changed across time points in both muscle types (**Figure 4**). At birth, nearly one-quarter of mitochondrial surface area was in direct contact with adjacent organelles in both muscle types (**Figure 4a,b,i**, 24.6±2.9 and 23.7±1.7% surface area contact per mitochondrion, n = 3 datasets each for P1 soleus and P1 gastrocnemius, respectively). However, by P14, membrane contact site abundance had dropped by nearly half and remained at that level throughout development (**Figure 4c-i**, 13.0±0.7, 11.9±1.3, 9.6±1.1, and 11.9±1.0% surface area contact per mitochondrion, n = 3 datasets each for P14 soleus, P14 gastrocnemius, P42 oxidative, and P42 glycolytic, respectively). These data suggest that mitochondria-organelle interactions may play a critical role in coordinating muscle cell assembly during the early postnatal period, while fewer interactions are needed to provide specialized support for muscle contraction in mature muscle cells.

## Discussion

By performing subcellular connectomic analyses of the nanoscale 3D mitochondria-organelle interactions within skeletal muscle cells across postnatal development, we demonstrate a physical reorganization of the cellular energy distribution system reflecting a functional transition away from cellular assembly and towards focused support of contractile function. Skeletal muscle in newborn mice were characterized by tortuous mitochondrial networks arranged in parallel to the contractile axis of the cell with frequent contact sites between mitochondria and lipid droplet or sarcoplasmic reticulum membranes. However, despite being placed directly between the myofibrils, the worm-like appearance of the neonatal mitochondria resulted in relatively less of the mitochondrial surface area being located directly adjacent to the contractile apparatus. Thus, the organization of the newborn muscle cell reflects an increased functional capacity for direct communication between organelles, lipid metabolism, and calcium signaling, the latter two of which have been suggested to play key roles in cellular assembly and development^42-46^. Conversely, regardless of whether the number of mitochondria increased (oxidative) or decreased (glycolytic) or whether mitochondria converted to grid-like (oxidative) or perpendicularly oriented (glycolytic) networks, interactions between mitochondria and other organelles fell nearly two-fold during postnatal development while the more linear mitochondrial segments in mature muscles formed tighter associations with the adjacent contractile structures. These data suggest that the mitochondrial outer membrane surface area in mature muscle is specialized to directly support the energetic requirements of muscle contractions independent of mitochondrial content or network configuration.

Individual mitochondria also appear capable of specialization relative to their adjacent neighbors within a mitochondrial network^10^. We show here that mitochondria connected to lipid droplets throughout postnatal skeletal muscle development are larger and more connected to adjacent mitochondria compared to non-lipid droplet connected mitochondria which have greater SA/V ratios. Thus, muscle mitochondria connected to lipid droplets have a greater structural capacity for energy conversion (i.e., greater internal volume) and rapid molecular transfer through mitochondrial networks (i.e., greater IMJ area) and less capacity for interactions with other cellular structures (i.e., lower available surface area relative to volume). This morphological distinction of lipid droplet connected mitochondria is consistent across postnatal development and has also been reported in mature heart muscle^10^ suggesting that lipid droplet connected mitochondria in all striated muscles may be functionally specialized to utilize the adjacent lipid fuel source and distribute the converted energy throughout the mitochondrial network to adjacent mitochondria which are structurally more suited for molecular distribution to other cellular components. However, while the relationship formed by membrane contact sites between mitochondria and lipid droplets appears consistent across striated muscles, a different type of functional specialization occurs in brown adipose tissue where lipid droplet connected mitochondria appear to be functionally specialized to support building lipid droplets rather than breaking them down for fuel^5,47^. As such, future work to gain a better understanding of the molecular nature of the tethers linking lipid droplets and mitochondria and how it may differ between muscle and brown fat cells may offer some insight into how certain tethers facilitate specific functions.

Contact sites between mitochondria and the endo/sarcoplasmic reticulum have been shown to play important roles in many cellular processes including mitochondrial dynamics^48,49^, calcium signaling^50,51^, lipid synthesis^50^, ubiquinone synthesis^52^, and cell death^53^. However, while the specific functional role of interactions between mitochondria and SR in skeletal muscle remains unclear^54^, the focus of much of the muscle work to date has revolved around the role of these organelle interactions in calcium signaling. Previous investigation into the postnatal relationship between mitochondria and SR/t-tubule triads found that the abundance of tethers linking mitochondria to these calcium release units (CRUs) was greater in mature than in two-week old mouse muscles^55^. Conversely, we found here that mitochondria-SR contact site abundance was greatest at birth and then fell significantly at two weeks of age and was maintained at the lower level until maturation. While the 3D nature of the current analyses versus the 2D nature of the previous study^22^ may play a role in this discrepancy, it is more likely due to the specificity of the previous analyses only to mitochondria-SR tethers located at CRUs whereas our analysis was not restricted to specific subcellular domains. The functional coupling of the SR and t-tubule triads likely specializes the calcium signaling in this region to support skeletal muscle excitation-contraction coupling. Thus, it is not surprising, and is consistent with the rest of this study, that mitochondrial interactions with CRUs would increase during the postnatal developmental reorganization period in order to more optimally support contraction in mature muscles. The current data combined with the previous study also suggest that it is the regions of the SR that are not part of the triads that account for the high abundance of mitochondria-SR contacts observed in the muscle at birth here. Indeed, in neonatal muscles, the SR is frequently associated with ribosomal granules^56^, which suggests that the SR has a greater capacity for classical endoplasmic reticulum functions such as protein synthesis^57,58^ at this time point and it is consistent with a role in cellular assembly. Additionally, calcium signaling may still take place in the non-triadic mitochondria-SR contacts observed in neonatal muscles, as calcium activity beyond its role in muscle contraction is known to be important for postnatal muscle development and cellular assembly^42-44^, and the abundance of the mitochondrial calcium uniporter is higher in both myotubes and neonatal muscles than in mature skeletal muscle^27,59^. Again, detailing the molecular nature of the tethers linking the SR and mitochondria in skeletal muscle and how it may differ between subcellular domains would provide key insight into the major molecular exchanges occurring at these specialized contact sites.

Building a skeletal muscle cell into a specialized contractile fiber requires the coordination of several different organelles, first to assemble and synthesize the structures needed, and then to reorganize the cell into the optimal configuration to meet the given contractile demands. During the neonatal phase in which cellular assembly is ongoing, the mitochondrial outer membrane maintains abundant contact with both the SR and large lipid droplets. In the dynamic phase of the transition towards functional specialization for contraction, the frequency of interactions between mitochondrial and SR membranes in the muscle cell decreases quickly, whereas the reduction in mitochondria-lipid droplet interactions drops more gradually. Finally, upon maturation, the mitochondrial outer membrane surface area becomes optimized to provide energetic support through molecular exchange with the surrounding contractile apparatus. Thus, these data reflect that, in addition to cellular organelle content and composition, organelle configuration is also highly tuned to the functional demands of the cell.

## Methods

### Mice

C57BL6/N mice (∼6-8 weeks old) were obtained from Taconic Biosciences (Rensselaer, NY) and were used as breeding pairs. Their offspring was randomly assigned as postnatal (P) 1, P14, and P42 group (n=3-4 per group). All pups were weaned at P21-P28, and both breeders and weaned animals accessed to food and water *ad libitum*. The vivarium was maintained on a 12:12 h light and dark cycle at 20-26 °C. Due to a technical limitation in using anogenital distance to ascertain sex in P1 pups, we randomly used male and female animals rather than separating experimental groups by sex. All animal procedures were approved by the National Heart, Lung, and Blood Institute Animal Care and Use Committee and performed in accordance with the guidelines described in the Animal Care and Welfare Act (7 USC 2142 § 13).

### Muscle Sample Preparation

As in a previous study^10^, both soleus (oxidative) and gastrocnemius (glycolytic) muscle fibers were carefully excised and fixed for FIB-SEM imaging acquisition. Briefly, skin-peeled hindlimbs were submerged in fixative solution (2% glutaraldehyde in 0.1M phosphate buffer, pH 7.2) for 30 minutes, while mice were under anesthetization with 2% isoflurane by nosecone. After one-hour incubation in standard fixative solution (2.5% glutaraldehyde, 1% formaldehyde, 0.12 M sodium cacodylate, pH 7.2–7.4), the excised tissues were subsequently post-fixed and *en bloc* stained (i.e., 2% aqueous osmium tetroxide incubation) following the published protocol with minor modifications^10^. After an overnight incubation in 1% uranyl acetate solution at 4 °C, the samples were incubated at 60 °C for 20 min with Walton’s lead aspartate (0.02 M lead nitrate, 0.03 M aspartic acid, pH 5.5) and were thoroughly washed with distilled H2O at room temperature. Afterwards, the samples were gradually dehydrated with ethanol and were then incubated in 25%, 50%, 75%, and 100% Embed 812 resin solutions for ∼36 hours. Then, the tissue samples were placed on ZEISS SEM specimen mounts (Electron Microscopy Sciences, #75225, USA) and were polymerized at 60 °C for 2-3 days. After the polymerization, the samples were cut and polished by Leica UCT Ultramicrotome (Leica Microsystems Inc., USA) that carried Trimtool 90 diamond knives (DiATOME, Switzerland).

### FIB-SEM imaging

The ZEISS crossbeam 540 (Gemini II) was used to collect FIB-SEM images at 10 nm voxel size with ZEISS Atlas 5 software (Carl Zeiss Microscopy GmbH, Jena, Germany). The FIB milling (10 nm thickness) was conducted at 30 keV, while maintaining a beam current at 2–2.5 nA. All collected micrographs were aligned with a proprietary algorithm and then exported as 8-bit TIFF files for further imaging analysis.

### Segmentation of cellular structures

All image processing was conducted on a desktop PC (New Tech Solutions, Fremont, CA) equipped with an Intel Xeon W-2155 (3.3 GHz processor, 10 cores/20 threads) and 256 GB RAM. As done previously^10^, Ilastik pixel classification software (Ilastik.org) was used for semi-automated image classification and segmentation of 3D structures of mitochondria and other subcellular organelles including lipid droplets, sarcoplasmic reticulum, and t-tubules. Also as done previously^10^, individual mitochondria were segmented using the Multicut module in Ilastik. All data were exported as a 32-bit HDF file for imaging analysis with ImageJ (National Institutes of Health, Bethesda, MD, ImageJ.net).

### Imaging Analysis

After loading with the HDF5 plugin, all HDF image datasets were processed to investigate both mitochondrial network configurations and individual mitochondrial structures. Following the established analytical pipeline^10^, several ImageJ plugins and analytical tools were used to examine mitochondrial networks (OrientationJ Distributions plugin), individual mitochondrial structures (ROIManager3D plugin), intermitochondrial junctions (Image Calculator tool), and mitochondrial spatial interactions with other subcellular components (Image Calculator tool and 3D Geometrical Measure tool). All 3D mitochondrial and subcellular components were extracted and visualized using 3D viewer and volume viewer on ImageJ or in Imaris.

### Statistical Analysis

Using Excel 2016 (Microsoft, Redmond, WA) and Prism 8 (GraphPad, San Diego, CA), we conducted all statistical analyses. Two-Way ANOVA was used to assess mean values of each dataset within and between groups (Fiber Type [Glycolytic, Oxidative] × Development [P1, P14, P42]). Tukey’s HSD *post hoc* tests were used for multiple comparisons and a statistical significance level was set at *P* < 0.05.

## Data and Code Availability

The raw FIB-SEM datasets for the present study have not been deposited in a public repository but are available from the corresponding author on reasonable request.

## Acknowledgements

This study was supported by the Division of Intramural Research of the National Heart Lung and Blood Institute and the Intramural Research Program of the National Institute of Arthritis and Musculoskeletal and Skin Diseases.

## Author Contributions

YK and BG prepped tissues for imaging. YK, EL, CKEB, and BG designed and EL and CKEB conducted imaging acquisitions. YK and BG designed and performed imaging analysis and made figures. YK and BG wrote the manuscript. YK, EL, CKEB, and BG edited and approved the manuscript.

## Declaration of Interests

The authors declare no competing interests.

## Notes

### Competing Interest Statement

The authors have declared no competing interest.

